# ^1^H, ^15^N and ^13^C sequence specific backbone assignment of the MAP Kinase Binding Domain of the Dual Specificity Phosphatase 1 and its interaction with the MAPK p38

**DOI:** 10.1101/2020.11.26.399634

**Authors:** Ganesan Senthil Kumar, Rebecca Page, Wolfgang Peti

## Abstract

The sequence-specific backbone assignment of the mitogen-activated protein kinase (MAPK) binding domain of the dual-specificity phosphatase 1 (DUSP1) has been accomplished using a uniformly [^13^C,^15^N]-labeled protein. These assignments will facilitate further studies of DUSP1 in the presence of inhibitors/ligands to target MAPK associated diseases and provide further insights into the function of dual-specificity phosphatase 1 in MAPK regulation.

## Biological context

Dual-specificity phosphatases (DUSPs; 25 members) belong to the family of protein tyrosine phosphatases, a subset of which dephosphorylate MAP kinases (MAP kinase phosphatases or MKPs) (Caunt and Keyse 2013) (Lawan et al. 2013). Timely and spatially accurate dephosphorylation of MAPKs by MKPs is critical for the regulation of MAPK signaling pathways. Hence, MKPs are promising candidates for manipulating MAPK-dependent immune responses to enhance or reduce their activity in cancers, infectious diseases, or inflammatory disorders (Jeffrey et al. 2007) (Ducruet et al. 2005) (Keyse 2008). MKPs harbor an N-terminal MAPK binding domain (MKBD) and a C-terminal catalytic domain (PTP) (Patterson et al. 2009). MKPs have similar PTP catalytic domains. However, they differ in their MKBDs, which enables specific interactions with different MAPKs (Patterson et al. 2009). Critically, the few reported structures of MKP MKBDs are structurally distinct; if and how this structural diversity enables specific MAPK binding has not been determined (Farooq et al. 2001) (Tao and Tong 2007) (Zhang et al. 2011). Hence, it is important to characterize the structure of MKPs to understand how they function, given the limited knowledge of known MKP structures.

DUSP1/MKP-1 was the first discovered MKP and clinical studies have shown that DUSP1 expression is correlated with multiple cancers (Charles et al. 1992) (Shen et al. 2016). For instance, DUSP1 is highly expressed in the malignant tissues of human breast cancer patients compared to non-malignant samples. Clinical studies have also shown that DUSP1 is a useful prognostic marker as its expression is correlated with cancer progression (Shen et al. 2017). The increased expression of DUSP1 correlated with reduced JNK activity, suggesting that therapies that reduce the expression or activity of DUSP1 might enable the expression of the pro-apoptotic signaling by JNK/p38 in malignant cells (Taylor et al. 2013). Interestingly, DUSP1 dephosphorylates all MAPKs (p38, JNK, and to a lesser extent ERK) (Patterson et al. 2009).

The key challenge of MAPK inhibitors is the broad expression profile and feedback loops of MAPKs resulting in a plethora of potential side effects. Hence, the modulation of DUSP activity is an alternative strategy for manipulating MAPK pathways in a cell-specific manner. However, the shallow active site (~ 6 Å compared to 9 Å for tyr-specific PTPs) and the hydrophilic nature of the catalytic domain presents challenges for developing drugs against DUSP activity (Tonks 2013). Hence, targeting the protein-protein interaction between the MKP MKBD and the MAPK is a promising alternative strategy for manipulating MKP activity and function. To this end, we initiated a solution NMR study of DUSP1 MKBD to study its interaction with MAP kinases. Herein we report the sequencespecific backbone resonance assignment of DUSP1 MKBD, which is the first step towards the characterization of their mode(s) of interaction. We also identified the interaction site of DUSP1 MKBD with p38.

## Methods and experiments

### Protein expression and purification

The coding sequence of the DUSP1 MAP kinase domain (MKBD, corresponding to residues 3-148) was sub cloned into RP1B (Peti and Page 2007). For protein expression, plasmid DNA was transformed into *E. coli* BL21 (DE3) RIL cells (Agilent). [^1^H, ^15^N, ^13^C]- or [^2^H,^15^N,^13^C]-labeled DUSP1 MKBD was achieved by growing cells in either H_2_O- or D_2_O-based M9 minimal medium containing selective antibiotics and 4 g/l [^13^C]/[^2^H,^13^C]-D-glucose and 1 g/L ^15^NH4Cl (CIL) as the sole carbon and nitrogen sources, respectively. Multiple rounds (0, 30%, 50%, 70%, and 100%) of D_2_O adaptation were necessary for high-yield protein expression (Peti and Page 2016). Expression was induced by the addition of 1 mM isopropylthio-*β*-D-galactoside (IPTG) when the optical density (OD_600_) reached 0.8-1.0. Induction proceeded overnight at 18 °C before harvesting the cells by centrifugation at 6000 *xg* (15 minutes, 4 °C). Cell pellets were stored at −80°C until purification. The expression and purification of human p38α (hereafter p38; residues 2-349) were carried out as previously described (Kumar et al. 2018).

Cell pellets were resuspended in lysis buffer (50 mM Tris-HCl pH 8.0, 500 mM NaCl, 5 mM imidazole, 0.1 % Triton X-100 containing EDTA-free protease inhibitor tablet [Roche]), lysed using high-pressure cell homogenization (Avestin C3 EmulsiFlex) and centrifuged (42,000 x*g*, 45 minutes, 4°C). The supernatant was filtered, loaded onto a HisTrap HP column (GE Healthcare) pre-equilibrated with Buffer A (50 mM Tris-HCl pH 8.0, 500 mM NaCl, 5 mM imidazole) and eluted using a linear gradient of Buffer B (50 mM Tris-HCl pH 8.0, 500 mM NaCl, 500 mM imidazole). Fractions containing the MKBD were pooled and dialyzed overnight with TEV protease in dialysis buffer (50 mM Tris pH 8.0, 500 mM NaCl) at 4°C. The next day, a ‘subtraction’ His-tag purification was performed to remove TEV and the cleaved His-tag. Final purification was achieved using size exclusion chromatography (SEC; Superdex 75 26/60 [GE Healthcare]) equilibrated in either NMR Buffer A (20 mM sodium phosphate pH 6.5, 100 mM NaCl, 0.5 mM TCEP), NMR Buffer B (50 mM HEPES pH 6.8, 150 mM NaCl, 5 mM DTT) or ITC Buffer (10 mM HEPES pH 7.5, 0.15M NaCl, 0.5 mM EDTA, 1 mM TCEP). The complex of [^2^H,^15^N,^13^C]-labeled DUSP1 MKBD and unlabeled p38 was generated by mixing equimolar ratios of the proteins followed by purification using SEC (Superdex 75 26/60, pre-equilibrated in NMR Buffer B).

### NMR spectroscopy

All NMR measurements were performed at 298 K on either a Bruker Avance II 500 MHz or 800 MHz spectrometer both equipped with TCI HCN Z-gradient cryoprobe. NMR samples were prepared in NMR buffer containing 10% (v/v) D_2_O. Sequence-specific ^1^H, ^15^N and ^13^C resonance assignment for DUSP1 MKBD (in NMR Buffer A; 0.4 mM) was obtained by analyzing 2D [^1^H,^15^N] HSQC, 3D HNCA, 3D HNCACB, 3D CBCA(CO)NH, 3D (H)CC(CO)NH (τ_m_=12 ms) and 3D HBHA(CO)NH spectra. 2D [^1^H, ^15^N] TROSY and a 3D HNCA-TROSY spectrum of the complex between unlabeled p38/[^2^H,^15^N,^13^C]-labeled DUSP1 MKBD (MW 56 kDa; NMR Buffer B: 0.5 mM) was used for the sequence-specific backbone assignment of DUSP1 MKBD in complex with p38. ^15^N[^1^H]-NOE (hetNOE) measurements were determined from a pair of interleaved spectra acquired with or without presaturation and a recycle delay of 5 seconds at 500 MHz ^1^H Larmor frequency. All NMR spectra were processed and analyzed using Topspin 2.1/3.0/3.1 (Bruker, Billerica, MA) or NMRPipe (Delaglio et al. 1995) and CARA (http://cara.nmr.ch) or NMRFAM-Sparky (Lee et al. 2015), respectively. Backbone amide chemical shift deviations were calculated using the formula: Δδ_av_=√(0.5 ((δ_HN,bound_-δ_HN,free_)^2^+ 0.04 δ_N,bound_-δ_N,free_)^2^)). Chemical shift indices (CSI) and secondary structure propensities (SSP) were calculated as previously described (Marsh et al. 2006) using the RefDB random coil database (Zhang et al. 2003).

### Isothermal Titration Calorimetry

ITC experiments were performed at 25°C using a VP-ITC microcalorimeter (Microcal Inc.). Titrant (10 μL per injection) was injected into the sample cell for 20 seconds with a 250 second interval between titrations to allow for complete equilibration and baseline recovery. 28 injections were delivered during each experiment and the solution in the sample cell was stirred at 307 rpm to ensure rapid mixing. To determine the thermodynamic parameters (ΔH, ΔS, ΔG) and binding constants (K), the DUSP1 MKBD was titrated into p38 and the data analyzed with a one-site binding model assuming a binding stoichiometry of 1:1 using NITPIC, SEDPHAT and GUSSI (Scheuermann and Brautigam 2015) (Zhao et al. 2015). All data were repeated in triplicate.

## Assignment and data deposition

DUSP1 belongs to the family of dual-specificity phosphatases and harbors two key domains, the MKBD (MAP kinase binding domain; also known as a rhodanase domain) and the catalytic domain (**Fig. 1A**). The [^1^H,^15^N]-HSQC spectrum of the DUSP1 MKBD (residues 3-148) shows that the backbone amide resonances are well dispersed, as expected for a well-folded protein (**Fig. 1B**). We obtained the backbone assignment for all DUSP1 MKBD residues, with the exception of K97. Backbone C^α^ was assigned for all the residues except the C-terminal proline (P148) and N-terminal glycine. Side-chain C^β^ resonance assignments are 99.2%, complete. **Fig. 1C** shows the secondary structure propensity (SSP) and chemical shift index (CSI) based on the ΔC^α^-ΔC^β^ chemical shift values, showing that DUSP1 MKBD has a mixed α-helical and β-sheet structure, which is typical of MKBDs of other dual-specificity phosphatases. Most importantly, the residues that constitute the kinase interaction motif (KIM; residues 50-66) shows higher α-helical propensities, as observed for the MKBDs of DUSP16 (PDBID 2VSW) and DUSP10 (Tao and Tong 2007). Furthermore, ^15^N[^1^H]-NOE (hetNOE) analysis showed that MKBD has very limited fast time scale (ps/ns) dynamics, consistent with a well-folded protein and is similar to what is observed for the DUSP16 MKBD (**Fig. 1D**) (Kumar et al. 2013). All chemical shifts were deposited in the BioMagResBank (http://www.bmrb.wisc.edu) under accession number 50574.

**Figure 1.**
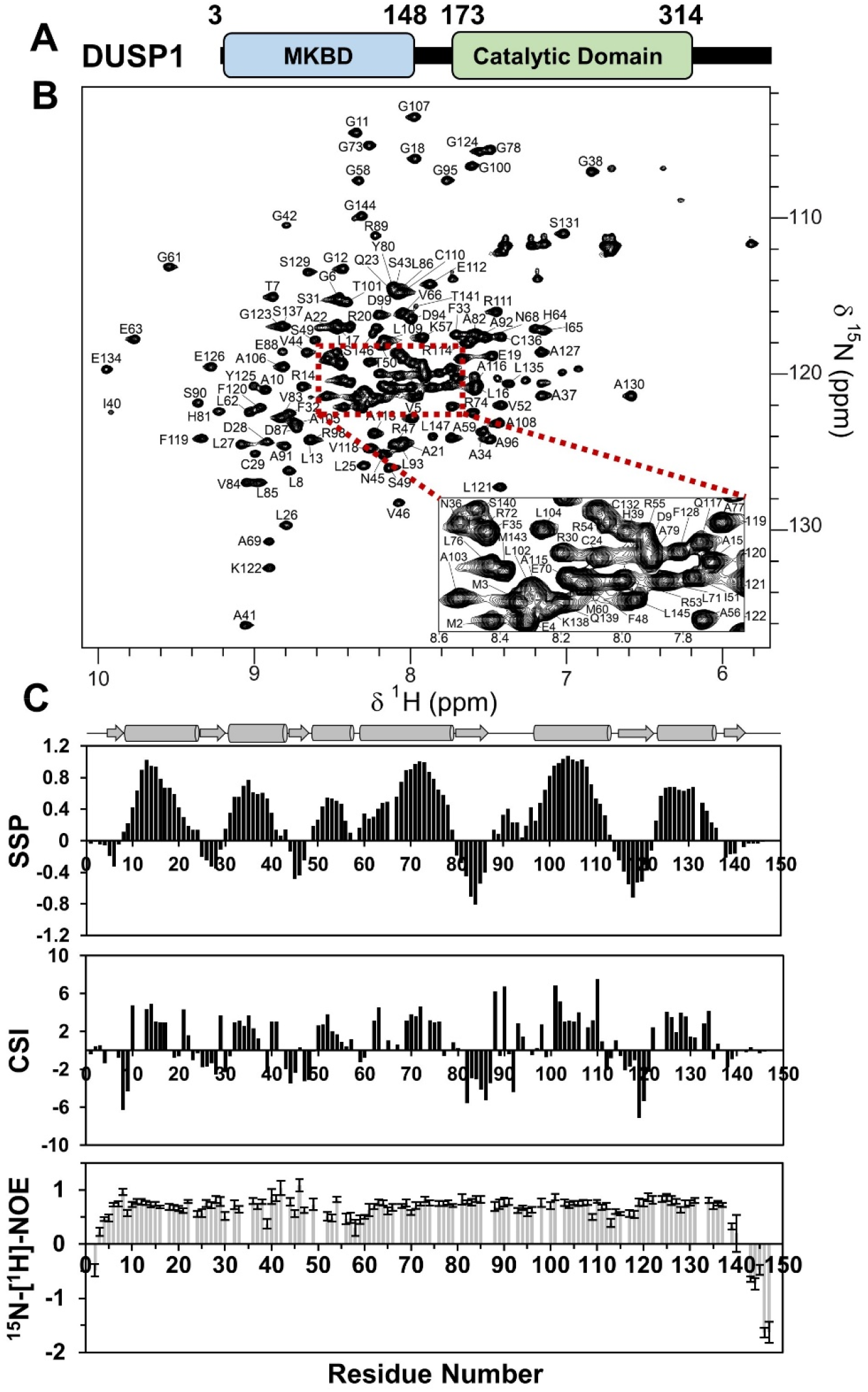
**(A)** DUSP1 has an N-terminal MAP kinase binding domain (MKBD, blue) and a C-terminal catalytic phosphatase domain (green). Domain boundary residues are shown. **(B)** Fully annotated 2D [^1^H,^15^N] HSQC spectrum of the DUSP1 MKBD. (**C**) Secondary structure propensities (SSP; *top panel*) and chemical shift index (CSI; *middle panel*) of the DUSP1 MKBD. Predicted secondary structural elements based on the SSP and CSI are also indicated. ^1^H[^15^N]-NOE of the DUSP1 MKBD (*bottom panel*), to show overall low fast-timescale flexibility of the DUSP1 MKBD, with the exception of the N- and C-termini.

## Protein Interaction Experiments

DUSP1 selectively inactivates JNK and p38 in response to external stimuli in the nucleus. To characterize the interaction of the DUSP1 MKBD with p38, we used isothermal titration calorimetry (ITC). ITC measurements showed that the DUSP1 MKBD binds strongly to p38, with a K_D_ of 445 ± 67 nM (**Fig. 2A; Table 1**). To identify the residues of DUSP1 MKBD the mediate p38 binding, we used solution NMR spectroscopy. The p38:DUSP1 MKBD complex was isolated using SEC. An overlay of the 2D [^1^H,^15^N] TROSY spectra of free- and p38 bound-DUSP1 MKBD showed large differences, making a direct assignment transfer impossible (**Fig. 2B**). Thus, to confirm the assignments, we completed the sequence-specific backbone assignment of the DUSP1 MKBD in complex with p38 by recording a 3D TROSY-HNCA spectrum. Comparison of the 2D [^1^H,^15^N] TROSY spectra of free and p38-bound DUSP1 MKBD shows CSPs of 23 peaks, 19 in fast exchange and 4 (T50, V52, M60 and G61) with linewidths broadened beyond detectability. All perturbed residues are localized to the Kinase interaction motif (KIM) binding sequence, confirming the importance of this motif for MAPK binding. The DUSP1 MKBD in the presence and absence of p38 showed similar secondary C^α^ chemical shifts indicating that the interaction with p38 does not result in any secondary structural element changes (**Fig. 2C**). Further NMR studies of the DUSP1 MKBD with inhibitors/ligands will aid the development of DUSP1 MKBD treatments for MAP kinase associated diseases.

**Figure 2.**
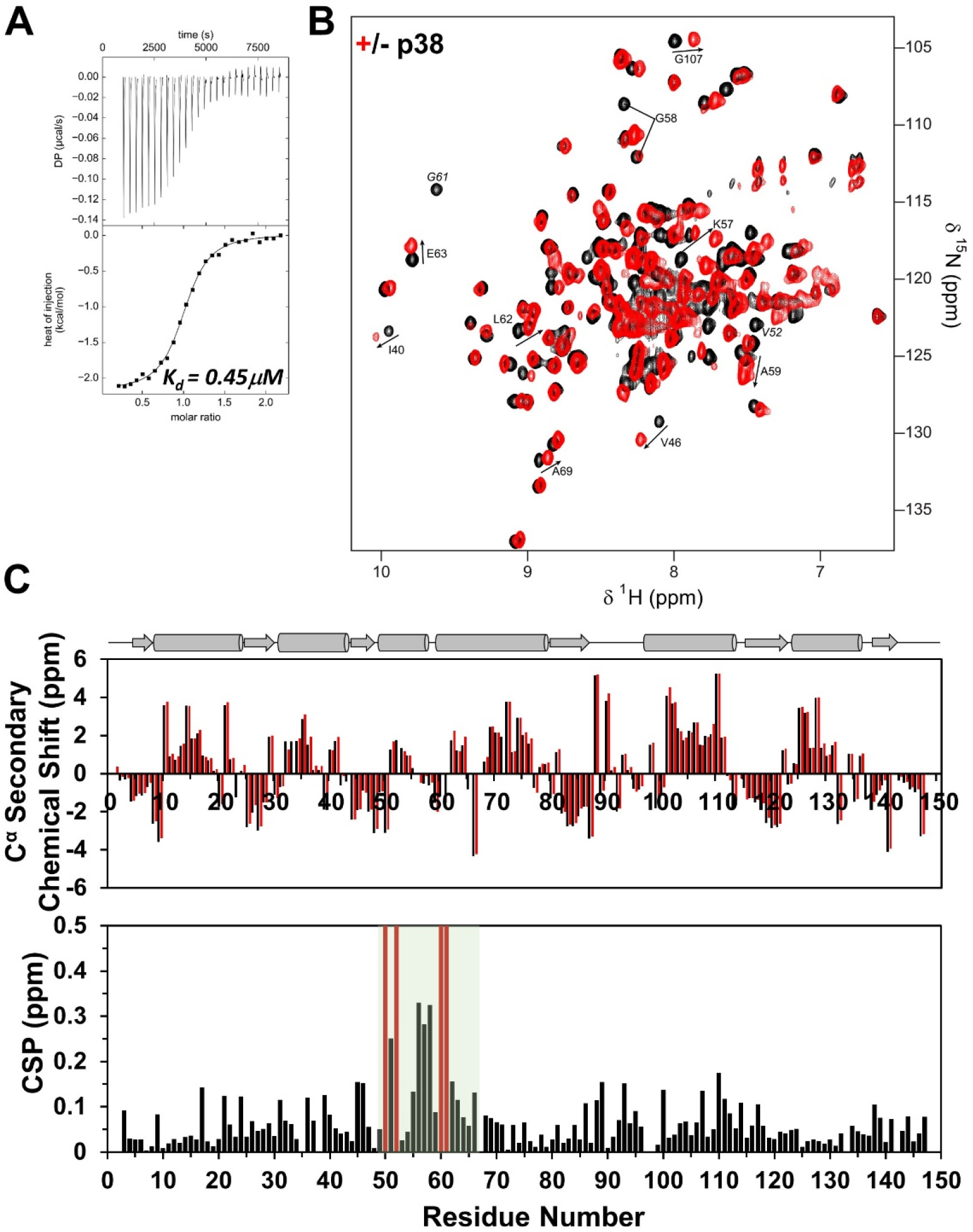
**(A)** Isothermal titration calorimetry of DUSP1 MKBD with p38; data recorded in triplicate. **(B)** Overlay of the 2D [^1^H,^15^N] TROSY spectrum of the (^2^H,^15^N)-DUSP1 MKBD in the presence (red) and absence (black) of p38. Residues with significant chemical shift changes are highlighted. The residues undergoing line broadening are annotated in italics. (**C**) C^α^ secondary shifts of DUSP-1 MKBD as a function of residue number in the presence (red) and absence (black) of p38 (*top panel*). Note that the secondary chemical shifts observed in the presence and absence of p38 are similar, showing that the secondary structural elements are conserved upon the complex formation. Chemical shift perturbations of DUSP-1 upon binding to p38 as function of residue number (*bottom panel*). The significant changes observed are localized to the KIM motif (highlighted by a green box). The residues line broadened beyond detection are indicated by red bars.

**Table 1.** Thermodynamic and dissociation constants for p38:DUSP1 MKBD derived from ITC experiments at 25°C.

| Peptide | $K_D$ (nM) | $\Delta G$ (kcal·mol<sup>-1</sup>) | $\Delta H$ (kcal·mol<sup>-1</sup>) | $T\Delta S$<br>(kcal·mol<sup>-1</sup>) |
| --- | --- | --- | --- | --- |
| p38:DUSP1 MKBD | 445 ± 67 | -8.7 ± 0.1 | -2.2 ± 0.2 | 6.4 ± 0.1 |

Experiment was conducted in triplicate.

## Acknowledgments

The authors thank Dr. Heiko Zettl for help at the early stages of the project. This research was supported by grant RSG-08-067-01-LIB from the American Cancer Society to R.P. and by R01GM100910 from the National Institute of Health to W.P. This research is based in part on data obtained at the Brown University Structural Biology Core Facility, which is supported by the Division of Biology and Medicine, Brown University. 800 MHz NMR data were recorded at Brandeis University.

## References

Caunt CJ, Keyse SM (2013) Dual-specificity MAP kinase phosphatases (MKPs): shaping the outcome of MAP kinase signalling. FEBS J 280:489–504. https://doi.org/10.1111/j.1742-4658.2012.08716.x

Charles CH, Abler AS, Lau LF (1992) cDNA sequence of a growth factor-inducible immediate early gene and characterization of its encoded protein. Oncogene 7:187–190

Delaglio F, Grzesiek S, Vuister GW, et al (1995) NMRPipe: a multidimensional spectral processing system based on UNIX pipes. J Biomol NMR 6:277–293. https://doi.org/10.1007/BF00197809

Ducruet AP, Vogt A, Wipf P, Lazo JS (2005) Dual specificity protein phosphatases: therapeutic targets for cancer and Alzheimer’s disease. Annu Rev Pharmacol Toxicol 45:725–750. https://doi.org/10.1146/annurev.pharmtox.45.120403.100040

Farooq A, Chaturvedi G, Mujtaba S, et al (2001) Solution structure of ERK2 binding domain of MAPK phosphatase MKP-3: structural insights into MKP-3 activation by ERK2. Mol Cell 7:387–399. https://doi.org/10.1016/s1097-2765(01)00186-1

Jeffrey KL, Camps M, Rommel C, Mackay CR (2007) Targeting dual-specificity phosphatases: manipulating MAP kinase signalling and immune responses. Nat Rev Drug Discov 6:391–403. https://doi.org/10.1038/nrd2289

Keyse SM (2008) Dual-specificity MAP kinase phosphatases (MKPs) and cancer. Cancer Metastasis Rev 27:253–261. https://doi.org/10.1007/s10555-008-9123-1

Kumar GS, Clarkson MW, Kunze MBA, et al (2018) Dynamic activation and regulation of the mitogen-activated protein kinase p38. Proc Natl Acad Sci U S A 115:4655–4660. https://doi.org/10.1073/pnas.1721441115

Kumar GS, Zettl H, Page R, Peti W (2013) Structural basis for the regulation of the mitogen-activated protein (MAP) kinase p38α by the dual specificity phosphatase 16 MAP kinase binding domain in solution. J Biol Chem 288:28347–28356. https://doi.org/10.1074/jbc.M113.499178

Lawan A, Shi H, Gatzke F, Bennett AM (2013) Diversity and specificity of the mitogen-activated protein kinase phosphatase-1 functions. Cell Mol Life Sci 70:223–237. https://doi.org/10.1007/s00018-012-1041-2

Lee W, Tonelli M, Markley JL (2015) NMRFAM-SPARKY: enhanced software for biomolecular NMR spectroscopy. Bioinformatics 31:1325–1327. https://doi.org/10.1093/bioinformatics/btu830

Marsh JA, Singh VK, Jia Z, Forman-Kay JD (2006) Sensitivity of secondary structure propensities to sequence differences between alpha- and gamma-synuclein: implications for fibrillation. Protein Sci 15:2795–2804. https://doi.org/10.1110/ps.062465306

Patterson KI, Brummer T, O’Brien PM, Daly RJ (2009) Dual-specificity phosphatases: critical regulators with diverse cellular targets. Biochem J 418:475–489. https://doi.org/10.1042/bj20082234

Peti W, Page R (2007) Strategies to maximize heterologous protein expression in Escherichia coli with minimal cost. Protein Expr Purif 51:1–10. https://doi.org/10.1016/j.pep.2006.06.024

Peti W, Page R (2016) NMR Spectroscopy to Study MAP Kinase Binding to MAP Kinase Phosphatases. Methods Mol Biol 1447:181–196. https://doi.org/10.1007/978-1-4939-3746-2_11

Scheuermann TH, Brautigam CA (2015) High-precision, automated integration of multiple isothermal titration calorimetric thermograms: new features of NITPIC. Methods 76:87–98. https://doi.org/10.1016/j.ymeth.2014.11.024

Shen J, Zhang Y, Yu H, et al (2016) Role of DUSP1/MKP1 in tumorigenesis, tumor progression and therapy. Cancer Med 5:2061–2068. https://doi.org/10.1002/cam4.772

Shen J, Zhou S, Shi L, et al (2017) DUSP1 inhibits cell proliferation, metastasis and invasion and angiogenesis in gallbladder cancer. Oncotarget 8:12133–12144. https://doi.org/10.18632/oncotarget.14815

Tao X, Tong L (2007) Crystal structure of the MAP kinase binding domain and the catalytic domain of human MKP5. Protein Sci 16:880–886. https://doi.org/10.1110/ps.062712807

Taylor DM, Moser R, Régulier E, et al (2013) MAP Kinase Phosphatase 1 (MKP-1/DUSP1) Is Neuroprotective in Huntington’s Disease via Additive Effects of JNK and p38 Inhibition. J Neurosci 33:2313–2325. https://doi.org/10.1523/JNEUROSCI.4965-11.2013

Tonks NK (2013) Protein Tyrosine Phosphatases: From Housekeeping Enzymes to Master-Regulators of Signal Transduction. FEBS J 280:346–378. https://doi.org/10.1111/febs.12077

Zhang H, Neal S, Wishart DS (2003) RefDB: a database of uniformly referenced protein chemical shifts. J Biomol NMR 25:173–195. https://doi.org/10.1023/a:1022836027055

Zhang Y-Y, Wu J-W, Wang Z-X (2011) A distinct interaction mode revealed by the crystal structure of the kinase p38α with the MAPK binding domain of the phosphatase MKP5. Sci Signal 4:ra88. https://doi.org/10.1126/scisignal.2002241

Zhao H, Piszczek G, Schuck P (2015) SEDPHAT--a platform for global ITC analysis and global multi-method analysis of molecular interactions. Methods 76:137–148. https://doi.org/10.1016/j.ymeth.2014.11.012

